# Re-examining supposed correlations between synonymous codon usage and protein bond angles in *E. coli*

**DOI:** 10.1101/2022.10.26.513858

**Authors:** Opetunde J. Akeju, Alexander L. Cope

## Abstract

A recent paper by Rosenberg et al. [2022] found a surprising correlation between synonymous codon usage and the dihedral bond angles of the resulting amino acid. However, their analysis did not account for the strongest known correlate of codon usage: gene expression. We applied the approach of Rosenberg et al. [2022] to simulated protein-coding sequences that (1) maintain the general relationship between codon usage and gene expression and (2) completely random codon usage. The analysis of the simulated data assuming a general relationship between gene expression and codon usage returned results remarkably similar to the real data. More concerning was the large number of significant results detected when sequences with random codon usage were analyzed. We believe that the specific results of Rosenberg et al. [2022] were confounded by the relationship between codon usage and gene expression, but also that their method is generally prone to detecting noise in protein bond angle distributions.

## Introduction

Codon usage bias (CUB), or the non-uniform usage of synonymous codons, has been observed across all domains of life [Plotkin and Kudla, 2011]. CUB is driven by a combination of both non-adaptive (e.g. mutation biases) and adaptive (e.g. natural selection for translation efficiency/accuracy) evolutionary forces [Ikemura, 1981, Gouy and Gautier, 1982, Bulmer, 1991, Drummond and Wilke, 2008, Shah and Gilchrist, 2011]. Empirical work has shown that changes to synonymous codon usage can affect co-translational protein folding and various computational studies have sought to determine if there is a general connection between codon usage and protein structure. Rosenberg et al. [2022] recently explored the relationship between synonymous codon usage and the dihedral bond angles that form a protein’s backbone. Using a method they developed to compare dihedral bond angle distributions across synonymous codons, they detected numerous statistically significant differences between the dihedral bond angle distributions of synonymous codons within the *E. coli* proteome. Although they qualify their results by stating that correlation does not imply causation, they hypothesize that differences in dihedral bond angle distributions between synonymous codons could be due to differences in elongation speeds between codons. This is evidenced, they claim, by a correlation between the differences in the dihedral bond angles and the elongation speed of the codons.

Gene expression is the strongest known correlate with CUB, with highly expressed genes biased towards codons that are translated more efficiently or accurately and lowly expressed genes biased towards codons favored by mutational biases Drummond and Wilke [2008], Shah and Gilchrist [2011]. A particularly popular population genetics model for studying codon usage patterns posits that codon usage is at selection-mutationdrift equilibrium (SMDE) [Li, 1987, Bulmer, 1991], and has been extended over the past decade to explicitly model the effects of gene expression on the strength of natural selection on codon usage [Shah and Gilchrist, 2011, Wallace et al., 2013, Gilchrist et al., 2015]. Although the work of Rosenberg et al. [2022] does not explicitly make statements about the nature of the evolution of CUB, the SMDE framework known as the Ribosomal Overhead Cost version of the Stochastic Evolutionary Model of Protein Production Rates (ROC-SEMPPR) provides an excellent tool for testing if the observations of Rosenberg et al. [2022] may be explained by the relationship between mutation bias, natural selection, and gene expression. We present results using simulated data suggesting the specific findings of Rosenberg et al. [2022] are likely spurious due to failure to control for these factors. We also present results that the approach developed by Rosenberg et al. [2022] is highly sensitive to noise in the data.

## Results

To test the robustness of the conclusions of Rosenberg et al. [2022], we re-create their analysis using two simulated datasets: (1) codon usage is consistent with the ROC-SEMPPR model (SMDE) and (2) synonymous codon usage is completely random (i.e. the probability of seeing a codon for an amino acid with *n*_*a*_ synonymous codons is 1*/n*_*a*_). A key point of these simulations is that they do not consider position-specific information. The locations of codons should be (mostly) randomized in both simulated datasets, eliminating any or most of the signal related to protein bond angles, should it exist.

The computational pipeline and the publicly-available data used to recreate their analysis were downloaded from the sources specified in [Rosenberg et al., 2022]. The pipeline was applied using the default settings to both the real and simulated protein-coding sequences. To analyze the simulated protein-coding sequences with the Rosenberg et al. [2022] pipeline, the represented protein-coding sequences were replaced with their simulated counterparts, retaining all other information as is from the real data, including the bond angles.

### The specific results of Rosenberg et al. [2022] were the result of gene-specific biases towards particular codons

We used the ROC-SEMPPR model to generate a simulated set of protein-coding sequences that accounts for intergenic variation in gene expression Gilchrist et al. 2015]. ROC-SEMPPR assumes codon usage is at SMDE and can estimate parameters reflecting natural selection on synonymous codon usage and mutation biases by accounting for variation in gene expression across protein-coding sequences. ROC-SEMPPR only considers how codon counts vary across protein-coding sequences and not within, making it blind to information relevant to position-specific effects, such as protein secondary structure and protein bond angles.

The *E. coli* K12 MG1655 protein-coding sequences were downloaded from NCBI-RefSeq (GCF 000005845.2). Rosenberg et al. [2022] did not restrict their analysis to a single strain of *E. coli*. Non-K12 MG1655 proteincoding sequences used by Rosenberg et al. [2022] were downloaded from the European Nucleotide Archive and appended to the *E. coli* K12 MG1655 protein-coding sequence FASTA file. We note that one proteincoding sequence (ENA AAL21040.1) used by Rosenberg et al. [2022] was annotated in the ENA as a *S. enterica* gene, but this was included for completeness. Excluding positions with missing codons in the real data, the simulated data contained 99% of the amino acid sites included in the real data. ROC-SEMPPR was fit to these protein-coding sequences using the AnaCoDa R package Landerer et al. 2018] to estimate parameters reflecting natural selection on codon usage Δ*η*, mutation bias Δ*M*, and an evolutionary average estimate gene expression *ϕ*. We note that the ROC-SEMPPR gene expression estimates were well-correlated with estimates taken from Ribo-seq data (Figure 1A, Spearman *R* = 0.48), suggesting an overall good model fit. Parameters estimated by ROC-SEMPPR were then used to simulate the codon usage of *E. coli* proteincoding sequences. Briefly, for each occurrence of an amino acid in a protein-coding sequence *g*, a synonymous codon was randomly sampled according to a multinomial distribution with the probability *p*_*i,g*_ of observing synonymous codon *i* (of *N* synonymous codons) determined by the expression level of the gene *ϕ*_*g*_ and the natural selection Δ*η* and mutation bias Δ*M* parameters estimated by ROC-SEMPPR for the relevant set of synonymous codons (see Equation 1 and see Gilchrist et al. 2015] for details).

**Figure 1:**
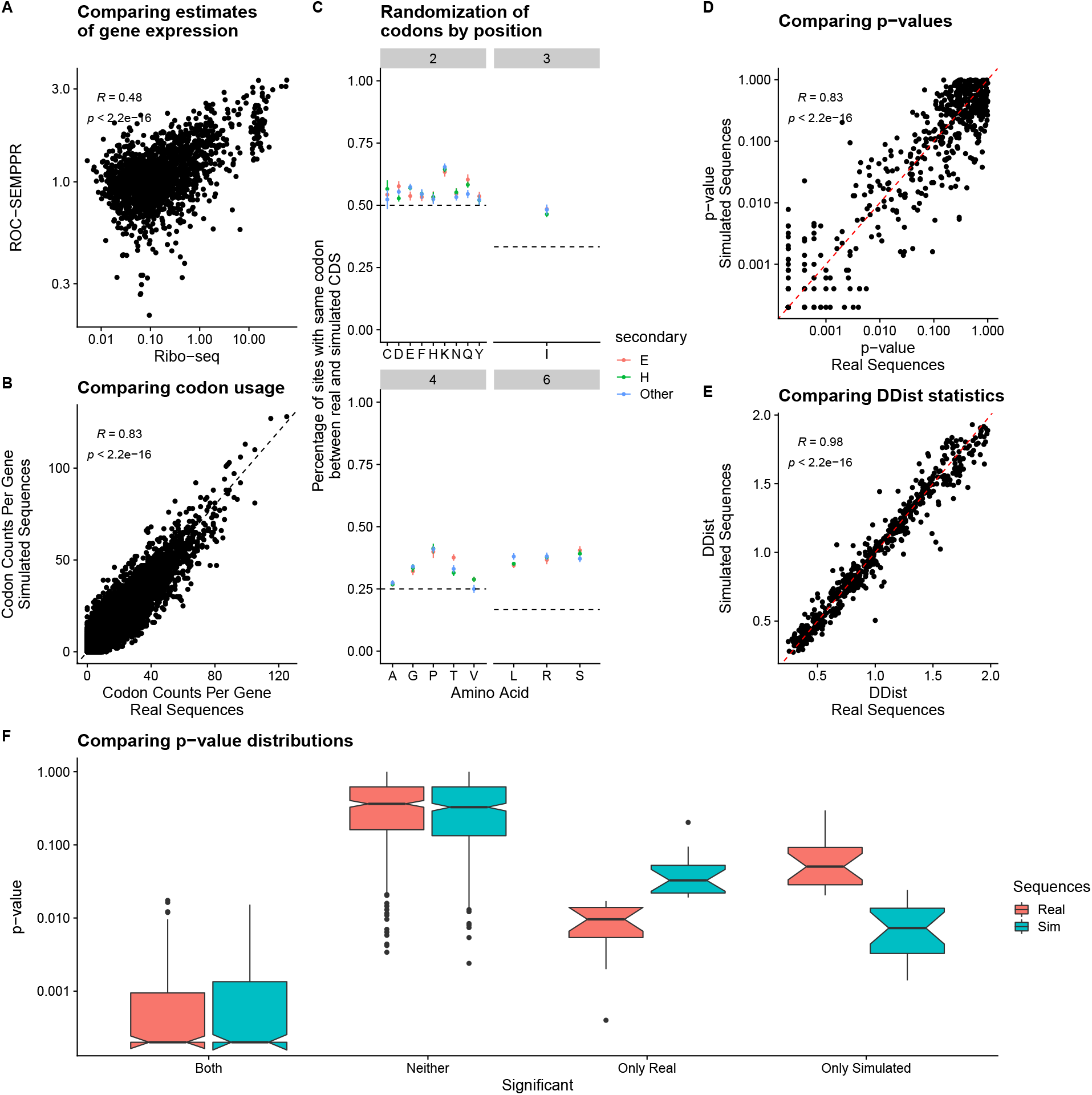
(A) Comparing estimates of ROC-SEMPPR predicted gene expression to Ribo-seq estimates taken from Mohammad et al. 2019]. (B) Comparing codon counts across all protein-coding sequences in the real and simulated data (ROC-SEMPPR). Each dot represents the number of occurrences of codon within a given gene (C) Percentage of amino acid sites that have the same codon at the same position in the same proteincoding sequence between the real and simulated data. The dashed line indicates the expectation under a completely random distribution. (D) Comparison of p-values estimated from the Rosenberg *et al*. pipeline applied to the real and simulated data. (E) Comparison of distribution distance statistics estimated from the Rosenberg *et al*. pipeline applied to the real and simulated data. (F) Comparing p-value distributions for synonymous codons that were significant in both, neither, or only one of the real and simulated data analyses.

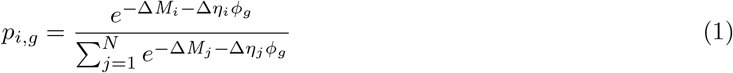

As expected, the real and simulated protein-coding sequences had similar codon counts per gene (Figure 1B, Spearman *R*_*s*_ = 0.83).

Although codon usage within the simulated protein-coding sequences is similar to the real sequences, the positions of synonymous codons within the sequences are effectively randomized (Figure 1C). For example, for the 4-codon amino acid valine (V), the percentage of times the same codon occurred at the same position within a protein-coding sequence between the real and simulated protein-coding sequences is approximately 25%. For many amino acids, there tends to be a small upward bias relative to the uniform expectation (Figure 1C, dashed lines). This is likely due to the fact that some protein-coding sequences will be strongly biased towards certain synonymous codons and that some codons are rarely used, such as the isoleucine codon ATA. Given the overall lack of agreement between the real and simulated sequences on a position-specific basis, the simulated sequences should contain little, if any, of the correlation between synonymous codon usage and protein bond angles, provided there is a general relationship between the two.

Overall, we found striking agreement between the analysis applied to the real and ROC-SEMPPR simulated sequences. 91% of the statistically significant synonymous codons using the real sequences were also found to be significant when using the simulated sequences, despite the overall randomization of codon usage by position. In agreement with this, the p-values and the distances between the bond angle distributions (DDist, to use the column name from pipeline output) between the two analyses were highly correlated (Figure 1D,E, Spearman *R*_*s*_ = 0.83 and Spearman *R*_*s*_ = 0.98, respectively). In cases where bond angle distributions for synonymous codons were only significant in one of the two datasets, the same codons tended to have a lower, but not significant, p-value in the other dataset (Figure 1F). Given that the simulated sequences under the ROC-SEMPPR model accurately reflect gene-specific codon usage patterns, but not position-specific codon usage patterns, it is likely the significant results obtained by Rosenberg et al. [2022] are heavily impacted by gene-specific biases. In the context of our results, the correlation between their distance metric and differences in codon-specific elongation speed (Figure 5 in Rosenberg et al. [2022]) suggests they are detecting signals related to selection for translation efficiency.

### Secondary structure frequencies and amino acid biases weakly vary with gene expression

What potential gene-specific biases could lead to significant differences in dihedral bond angle distributions between synonymous codons? A key result from Rosenberg et al. [2022] was that differences in bond angle distributions were present in *β*-sheets and not *α*-helices. We observe a weak correlation between the frequency of *β*-sheets in a gene and gene expression (Figure 2(right), Spearman correlation *R*_*s*_ = *−*0.14, *p* = 0.002). The correlation between gene expression and the frequency of *α*-helices in a gene is not statistically significant (Figure 2(left), Spearman correlation *R*_*s*_ = 0.078, *p* = 0.085). Amino acid usage is also known to correlate with gene expression in *E. coli* [Akashi and Gojobori, 2002]. Based on a correspondence analysis of amino acid usage across genes, we observe that gene expression is also weakly correlated with the second, third, and fourth principal components (Spearman correlation |*R*_*s*_| *<* 0.21, *p <* 0.006 in all cases), which in total “explain” 27.9% of the variation in amino acid usage across genes. The weak correlations of amino acid usage and secondary structure with gene expression may contribute to some of the apparent differences in the bond angle distributions between synonymous codons by introducing biases and/or further confounding comparisons between synonymous codons.

**Figure 2:**
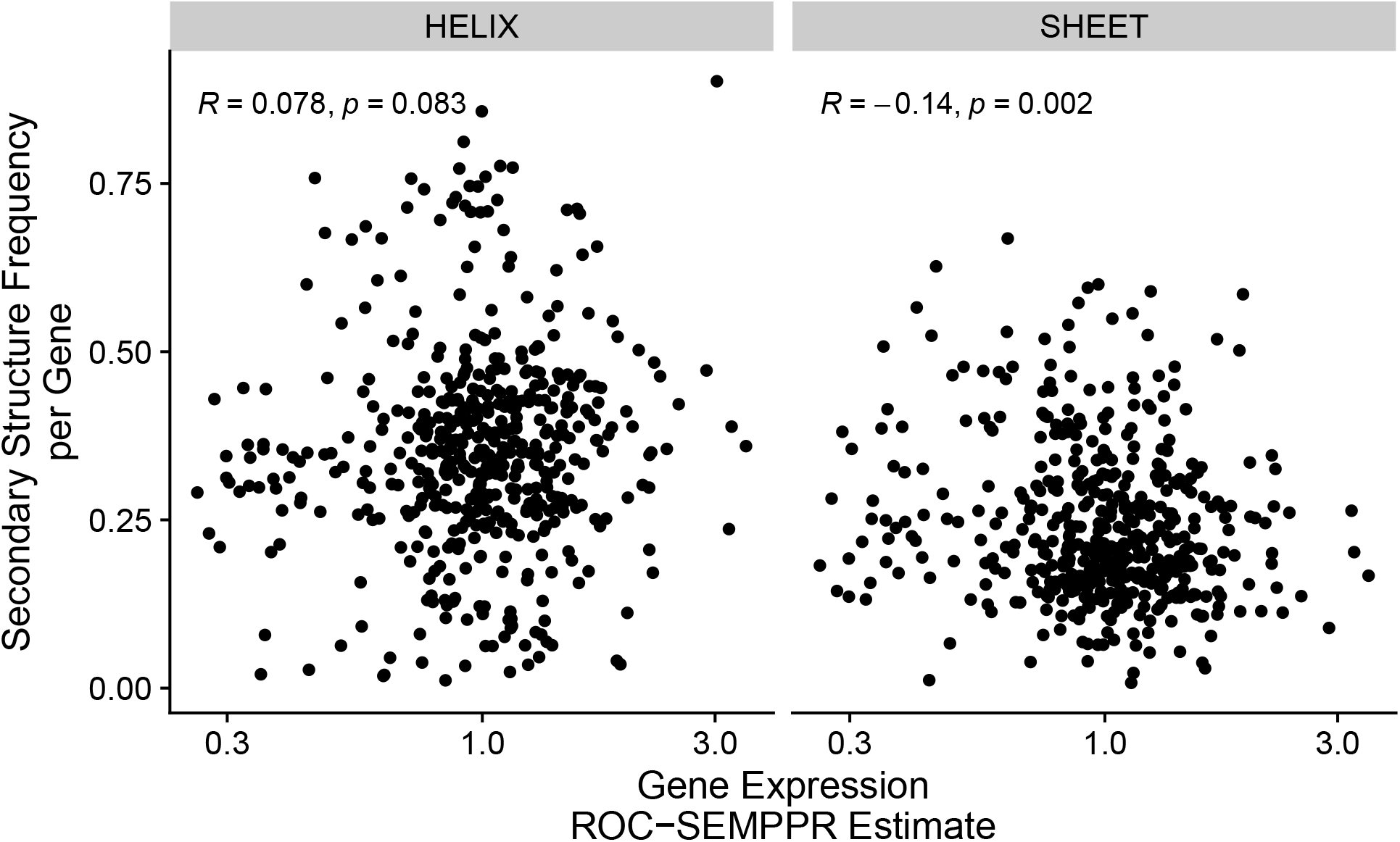
Correlation between gene expression estimate from ROC-SEMPPR *ϕ* and the frequency of each major secondary structure in each gene.

### The method of Rosenberg et al. [2022] is prone to attributing noise to signal

As an additional test, we generated a set of simulated protein-coding sequences in which synonymous codon usage was completely random. Again, one would expect there to be no signal relating codon usage to protein bond angles in this dataset. However, 78 pairs of synonymous codons were found to be significantly different in *β*-sheets in the randomly simulated protein-coding sequences compared to 58 in the real data. No synonymous codons were significantly different for *α*-helices in either dataset.

The striking number of statistically significant results in *β*-sheets under completely random codon usage suggests the method developed by Rosenberg et al. [2022] conflates noise in dihedral bond angles with signal, potentially due to over-fitting of the kernel density to the bond angle distributions. As an additional control, Rosenberg et al. [2022] applied their statistical test to compare the bond angle distributions of each codon to itself. As expected, a codon’s bond angle distribution is never significantly different from itself. However, there are still large variations in the calculated test statistics (the distances between the bond angle distributions) for each codon compared to itself, which reflects the degree of random noise in the protein bond angle distributions for a codon, as none of these distributions are truly different. We tested if amino acids that tended to show greater distances between bond angle distributions within codons also showed greater distances between synonymous codons within each secondary structure group. We observed a strong correlation between the mean test statistics within codons and between codons when using the real protein-coding sequences in all 4 classes of protein secondary structure (Figure 3). Perhaps more telling, we found that these correlations also exist in the analysis of the simulated data with completely random codon usage, potentially explaining why significant results were found where none should exist (Supplementary Figure S1). The strong correlations observed here suggest the Rosenberg et al. [2022] method is primarily detecting random noise.

**Figure 3:**
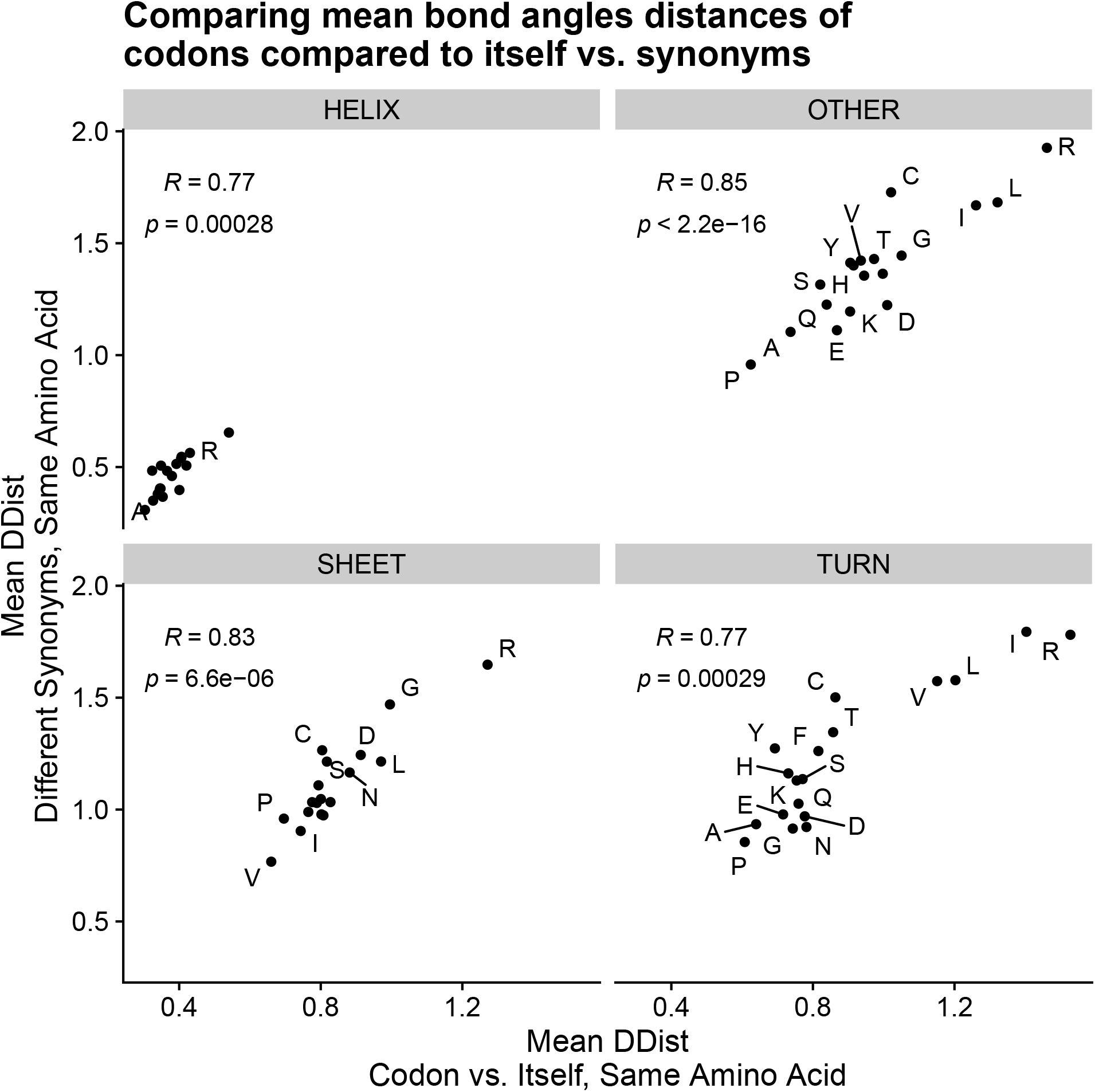
Comparison of the mean distances between bond angle distributions (the test statistic of Rosenberg et al. [2022]) when a codon is compared to itself vs. its synonyms for each amino acid and secondary structure. These are based on the analysis of the real protein-coding sequences. Note that the distances in bond angle distributions when comparing a codon with itself represent noise, as the underlying distributions are actually the same.

## Conclusion

Much like Rosenberg et al. [2022] could not definitively say if synonymous codon usage determined dihedral bond angles or vice versa, we cannot definitively say there is no relationship between synonymous codon usage and dihedral bond angles. Such a relationship would be truly remarkable: effects at the A-site of the ribosome would need to be carried over through the ribosome tunnel (approximately 35-40 amino acids long) and any subsequent co-translational or post-translational protein folding. As is, we remain highly skeptical.

Our results strongly suggest that the observed differences in bond angle distributions between codons are substantially impacted by the correlations between codon usage and gene expression. We find some evidence that secondary structure frequencies and amino acid usage vary as a function of gene expression; however, these correlations appear to be relatively weak. More concerning is that the approach developed by Rosenberg et al. [2022] is highly-prone to detecting random noise in bond angle distributions. Even when codon usage is completely random, we still observe numerous statistically significant results in *β*-sheets, but none in *α*-helices, consistent with analysis of the real data and data simulated under the ROC-SEMPPR model. Thus, while we believe the specific results of Rosenberg et al. [2022] were due to the relationship between codon usage and gene expression, the method itself appears to be generally unreliable. The relationship between codon usage, protein structure, and the protein folding process remains an exciting area of research, particularly as advances in molecular and structural biology allow for more detailed investigations and experiments. Further work is needed, particularly in the development of robust computational methods, in order to ascertain if a mechanistic relationship between protein bond angles and codon usage exists.

### Data availability

Relevant data and scripts can be found at https://github.com/acope3/Codon_usage_prot_structure_angles.

## Contributions

O.J.A. contributed to the analysis, and the writing and editing of the manuscript. A.L.C. conceptualized the project, contributed to the analysis, and helped write and edit the manuscript.

## Acknowledgments

We would like to thank Premal Shah, Michael Gilchrist, Matthew Pennell, and Joshua Schraiber for their helpful comments during the writing of this manuscript.

## Supplemental

**Figure S1:**
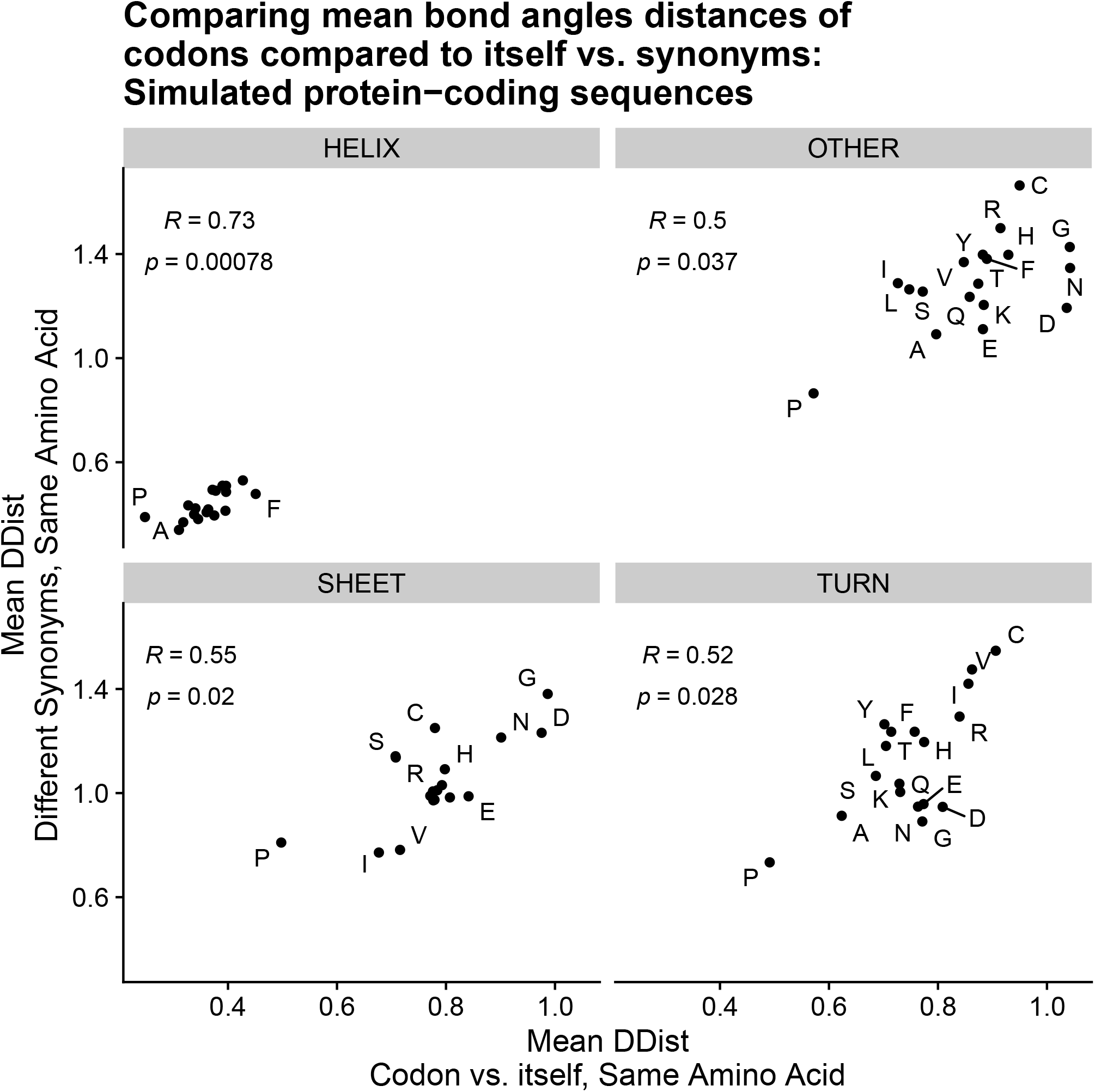
Comparison of the mean distances between bond angle distributions (the test statistic of Rosenberg et al. [2022]) when a codon is compared to itself vs. its synonyms for each amino acid and secondary structure. These are based on the analysis of the randomly simulated protein-coding sequences. Note that the distances in bond angle distributions when comparing a codon with itself represent noise, as the underlying distributions are actually the same.

